# GAD: a Python script for dividing genome annotation files into feature-based files

**DOI:** 10.1101/815860

**Authors:** Ahmed Karam, Norhan Yasser

## Abstract

Nowadays, manipulating and analyzing publicly available genomic datasets become a daily task in bioinformatics and genomics laboratories. The release of several genome sequencing projects prompts bioinformaticians to develop automated scripts and pipelines which analyze genomic datasets in particular gene annotation pipelines. Handling genome annotation files with fully-featured programs used by non-developers is necessary, furthermore, accelerating genomic data analysis with a focus on diminishing the genome annotation and sequence files based on specific features is required. Consequently, to extract genome features from GTF or GFF3 in a precise manner, GAD script (https://github.com/bio-projects/GAD) provides a simple graphical user interface which interpreted by all python versions installed in different operating systems. GAD script contains unique entry widgets which are capable to analyze multiple genome sequence and annotation files by a click. With highly influential coded functions, genome features such upstream genes, downstream genes, intergenic regions, genes, transcripts, exons, introns, coding sequences, five prime untranslated regions, and three prime untranslated regions and other ambiguous sequence ontology terms will be extracted. GAD script outputs the results in diverse file formats such as BED, GTF/GFF3 and FASTA files which supported by other bioinformatics programs. Our script could be incorporated into various pipelines in all genomics laboratories with the aim of accelerating data analysis.

## Introduction

Genome annotations are crucial for defining, accessing and extracting genomic parts with the standard set of definitions and terms (Eilbeck et al. 2005). GFF is a format that frequently used for storing genome annotation data. GFF2 (General Feature Format version 2) (http://gmod.org/wiki/GFF2) considered as a parent of all GFF file versions but it has several weak points, thence, GFF3 file (Generic Feature Format version 3) (http://gmod.org/wiki/GFF3) generated to make genomic features fully informative and well represented. In addition to, GTF file (Gene Transfer Format) (http://mblab.wustl.edu/GTF22.html) which is another genome annotation format vastly used alongside with GFF3 but governs different structures. Both GFF3 and GTF are nine-column tab-delimited files containing genomic feature coordinates per line.

There were several databases store genome sequences with their annotation files such as WormBase (Harris et al. 2009), FlyBase (Tweedie et al. 2008), The Arabidopsis Information Resource (Lamesch et al. 2011), Saccharomyces Genome Database (Cherry et al. 1998), Pseudomonas Genome DB (Winsor et al. 2010), National Center for Biotechnology Information (NCBI Resource Coordinators 2012) and Ensembl (Zerbino et al. 2017). Each of the above databases utilizes different approaches to annotate their genomes resulting in different structures of annotation file (Eilbeck et al. 2005). Nowadays, the Ensembl project processes publicly available data with automated annotation pipelines and storing the results in accessible databases with no restrictions (Zerbino et al. 2017).

The development of programs that have the potency to parse annotation files is strongly required to minimize the analysis efforts. Several programs handle annotation files such as BEDtools (Quinlan and Hall 2010), gffread (Trapnell et al. 2010), GFF-Ex (Rastogi and Gupta 2014) and gff2sequence (Camiolo and Porceddu 2013). As well, the galaxy project (Afgan et al. 2018) comprises a lot of programs that manipulate annotation files. Regrettably, no program found to deal with all categories of annotation files, genome features, devices, operating systems, and users. This encouraged us to fill the gap with our knowledge in the Python programming language; we release GAD (Genome Annotation Divider) script with simple, dynamic and controlled GUI (Graphical User Interface) for extracting genomic features from any GFF3, GTF file formats. GAD script interpreted by both python versions two and three installed on any operating systems and platforms. GAD script generates multiple file types which in turn reduce genome analysis efforts and resources.

## Implementation

The main algorithm of GAD script was coded in python programming language version 2 and interpreted by python version 3. The GUI was dynamically constructed using standard python GUI package (Tkinter), a GUI toolkit, which is found by default in the main python package.

## Results and Discussions

We developed GAD script (https://github.com/bio-projects/GAD) with simple, dynamic and controlled GUI for dividing annotation files into feature-based files. The single file input section provides simple widgets for uploading single GTF or GFF3 file format as a required entry and single FASTA file as an optional entry, the multiple files section having an algorithm for handling multiple annotation and sequence files with the same name in pairs form as well as sequence files optional to be uploaded, which we called pairing system.

All submitted files are parsed for extracting all required features; the select feature(s) section lets the user mark upon at least one interesting feature. GAD script includes a group of functions for extracting genomic regions such as intergenic regions, genes, mRNAs, exons, introns, CDSs, 5 prime UTRs, and 3 prime UTRs. Also, up- and down-stream could be extracted separately with special functions that require the fragment length to be generated in fixed or flexible length. Other ambiguous features also will be extracted by entering biotypes in a comma-separated style. Generated information is held to the unique output section, GAD script generates three output formats if marked for each feature, BED file is a lightweight file format containing specified genomic feature coordinates, in addition to annotation input file is curtailed to the same input format with required feature, and multiple FASTA files containing sequences corresponding to marked feature coordinates if nucleotide sequence in FASTA file is submitted.

Unlimited sequence and annotation files could be analyzed by one click with a unique pairing system currently exist which allows analyzing large datasets instantly. Six different genome sizes ranged from ~100 to ~15000 Mbp were downloaded as chromosome GFF3 and FASTA files from Ensembl plants (http://plants.ensembl.org/index.html) and analyzed (Table 1) by GAD script pairing system.

**Table 1:** Computer-clock time for analyzing genomes in the form of chromosomes by GAD pairing system

| Organism | Genome size (bp) | Time (Minutes) |
| --- | --- | --- |
| *Arabidopsis thaliana* | 135,670,229 | 1 |
| *Gossypium raimondii* | 747,964,930 | 2 |
| *Hordeum vulgare* | 8,059,674,078 | 13 |
| *Triticum aestivum* | 14,547,261,565 | 31 |
| *Triticum dicoccoides* | 10,494,678,545 | 27 |
| *Zea mays* | 2,104,350,183 | 6 |

The GAD pairing system provides the capability of using computers with low memory capacity in large genome analysis by a click. The chromosome file of a large genome utilization minimizes memory usage; on the other hand, the selection of multiple files is crucial for decreasing user efforts. For example, 4 GiB of memory is insufficient for analyzing *Triticum aestivum* genome (14 Gbps) by the gff2sequense and GFF-Ex programs; alternatively, chromosomes will be entered one by one 21 times to complete genome analysis. Thus, the GAD pairing system gives a shad of light for integrating old-computers in genome data analysis.

Furthermore, to prove the efficacy of our GAD script, Ensembl database (http://www.ensembl.org) was used as a standard to construct fair comparison, we downloaded GTF, GFF3 and FASTA files of four different genomes from different kingdoms (Supplementary Material) and analyze them with GAD script compared with the most utilized and similar program gff2sequence (Camiolo and Porceddu 2013). The comparison illustrates (Supplementary Material) the superiority of GAD script in several advantages. For example, GAD script identified 36845 genes in the *Oryza sativa* GFF3 file compared with 35825 genes with 6553 duplicated genes resulted from gff2sequence. The differences are due to gff2sequence incapable to identify all SO terms found in the GFF3 file. In the same way, the identified SO terms of genes influence the results of transcripts, exons, introns, upstream genes, downstream genes, and intergenic regions. On the other hand, 5` and 3` UTRs and CDSs carry only one SO term, gff2squence was differing in the results of 5` and 3` UTRs and CDSs because it concatenates grouped features into one sequence, thus, they were similar in both GAD and gff2sequence. Overlapped genes also have an impact on the final outcome, so, results of GAD script in upstream genes, downstream genes, and intergenic regions is superior compared with gff2sequence. GAD script found ambiguous SO terms (pre_miRNA; chromosome; supercontig; biological_region; SRP_RNA; RNase_MRP_RNA) which may be gene or transcript definitions, GAD lets the user inserts any interesting ambiguous SO term to be integrated into gene or transcript processing or to be extracted as an independent feature file. As well as GAD script generates the output in BED, GTF and/or GFF3 files. The lightweight of the GUI reflects the major advantage of GAD script, the memory usage of GAD GUI is very low (~20 MB) and thus saves space in memory for receiving as much as possible large files.

All analysis was performed using Lubuntu 18.04 LTS (Linux distribution) with CPU running on 2.40GHz and 4 GB of memory.

## Conclusion

We emerge GAD script a fully-featured GUI known up to now for extracting genomic features from both GTF and GFF3 files. Pairing system prevents the conflict of large data with weak resources. GAD could be involved in several data analysis (e.g., finding genes, SNPs (single nucleotide polymorphisms) within up- and down-stream, miRNA motifs in UTRs, etc.). Also, combining lightweight GAD outputs (e.g., BED, GFF3, etc.) with a web-based platform such as galaxy will reduce handling potential of large files and accelerating genomic research. GAD pairing system gives a shad of light for integrating old-computers in genome data analysis.

## Acknowledgments

Special thanks to Ahmed’s and Norhan’s families for their continuous support.

