## Supplementary Material for "GAD: a Python script for dividing genome annotation files into feature-based files"

**Title:**

**The name(s) of the author(s):**

Norhan Yasser^1^ and Ahmed Karam^2,*^

**The affiliation(s) and address(es) of the author(s):**

^1^Department of Biotechnology, Faculty of Science, Cairo University, Giza, Egypt.

^2^Department of Genetics, Faculty of Agriculture, Cairo University, Giza, Egypt.

**The e-mail address, and telephone number(s) of the corresponding author:**

**List of FTP links used in Table 1:**

Arabidopsis thaliana

GFF3: <ftp://ftp.ensemblgenomes.org/pub/release-42/plants/gff3/arabidopsis_thaliana>

FASTA: <ftp://ftp.ensemblgenomes.org/pub/release-42/plants/fasta/arabidopsis_thaliana/dna/>

Gossypium raimondii

GFF3: <ftp://ftp.ensemblgenomes.org/pub/release-42/plants/gff3/gossypium_raimondii>

FASTA: <ftp://ftp.ensemblgenomes.org/pub/release-42/plants/fasta/gossypium_raimondii/dna/>

Hordeum vulgare

GFF3: <ftp://ftp.ensemblgenomes.org/pub/release-42/plants/gff3/hordeum_vulgare>

FASTA: <ftp://ftp.ensemblgenomes.org/pub/release-42/plants/fasta/hordeum_vulgare/dna/>

Triticum aestivum

GFF3: <ftp://ftp.ensemblgenomes.org/pub/release-42/plants/gff3/triticum_aestivum>

FASTA: <ftp://ftp.ensemblgenomes.org/pub/release-42/plants/fasta/triticum_aestivum/dna/>

Triticum dicoccoides

GFF3: <ftp://ftp.ensemblgenomes.org/pub/release-42/plants/gff3/triticum_dicoccoides>

FASTA: <ftp://ftp.ensemblgenomes.org/pub/release-42/plants/fasta/triticum_dicoccoides/dna/>

Zea mays

GFF3: <ftp://ftp.ensemblgenomes.org/pub/release-42/plants/gff3/zea_mays>

FASTA: <ftp://ftp.ensemblgenomes.org/pub/release-42/plants/fasta/zea_mays/dna/>

**FTP Links used in the comparison between GAD and gff2sequence (Table 2 and Table 3):**

Drosophila melanogaster

GFF3: <ftp://ftp.ensemblgenomes.org/pub/release-42/metazoa/gff3/drosophila_melanogaster>

GTF: <ftp://ftp.ensemblgenomes.org/pub/release-42/metazoa/gtf/drosophila_melanogaster>

FASTA: <ftp://ftp.ensemblgenomes.org/pub/release-42/metazoa/fasta/drosophila_melanogaster/dna/>

Oryza sativa

GFF3: <ftp://ftp.ensemblgenomes.org/pub/release-42/plants/gff3/oryza_sativa>

GTF: <ftp://ftp.ensemblgenomes.org/pub/release-42/plants/gtf/oryza_sativa>

FASTA: <ftp://ftp.ensemblgenomes.org/pub/release-42/plants/fasta/oryza_sativa/dna/>

Plasmodium malariae

GFF3: <ftp://ftp.ensemblgenomes.org/pub/release-42/protists/gff3/protists_alveolata1_collection/plasmodium_malariae>

GTF: <ftp://ftp.ensemblgenomes.org/pub/release-42/protists/gtf/protists_alveolata1_collection/plasmodium_malariae>

FASTA: <ftp://ftp.ensemblgenomes.org/pub/release-42/protists/fasta/protists_alveolata1_collection/plasmodium_malariae/dna/>

Saccharomyces cerevisiae

GFF3: <ftp://ftp.ensemblgenomes.org/pub/release-42/fungi/gff3/saccharomyces_cerevisiae>

GTF: <ftp://ftp.ensemblgenomes.org/pub/release-42/fungi/gtf/saccharomyces_cerevisiae>

FASTA: <ftp://ftp.ensemblgenomes.org/pub/release-42/fungi/fasta/saccharomyces_cerevisiae/dna/>

Table 2: comparison between GAD and gff2sequence using GFF3 and FASTA files as inputs.

| Organism | *Drosophila melanogaster* | | *Oryza sativa* | | | | *Plasmodium malariae* | | | *Saccharomyces cerevisiae* | |
| --- | --- | --- | --- | --- | --- | --- | --- | --- | --- | --- | --- |
| Program/script | GAD | gff2sequence | | GAD | gff2sequence | GAD | | gff2sequence | GAD | | gff2sequence |
| Genes | 17753 | 13931 | | 36845 | 35825 | 6507 | | 6342 | 7036 | | 6599 |
| Gene SO names | gene;  ncRNA_gene;  pseudogene; | gene | | gene;  ncRNA_gene;  pseudogene; | gene | gene;  ncRNA_gene;  pseudogene; | | gene | gene;  ncRNA_gene;  pseudogene; | | gene |
| Transcripts | 34543 | 30504 | | 43298 | 42378 | 6507 | | 6342 | 7036 | | 6599 |
| Transcript SO names | mRNA;  ncRNA;  pseudogenic_transcript;  rRNA;  snRNA;  snoRNA;  tRNA; | mRNA | | lnc_RNA;  mRNA;  pseudogenic_transcript;  rRNA;  snRNA;  snoRNA;  tRNA; | mRNA | mRNA;  pseudogenic_transcript;  tRNA; | | mRNA | mRNA;  ncRNA;  pseudogenic_transcript;  rRNA;  snRNA;  snoRNA;  tRNA; | | mRNA |
| Up-stream | 17753 | 13928 | | 36845 | 35823 | 6507 | | - | 7036 | | 6574 |
| Dwon-stream | 17753 | 13929 | | 36845 | 35821 | 6507 | | - | 7036 | | 5477 |
| Intergenic regions | 18982 | 9354 | | 37125 | 30983 | 10314 | | 2054 | 7196 | | 5936 |
| 5`UTRs | 46335 | 29884 | | 45538 | 34863 | 337 | | 321 | 4 | | 4 |
| 3`UTRs | 33907 | 30490 | | 51046 | 35530 | - | | - | - | | - |
| Introns | 152724 | 150627 | | 149095 | 149079 | 7568 | | 7189 | 380 | | 313 |
| Exons | 187526 | 181131 | | 192493 | 191452 | 14075 | | 13532 | 7507 | | 6913 |
| CDSs | 160973 | 30504 | | 165077 | 42373 | 13532 | | 6342 | 6913 | | 6599 |
| Other feature(s) SO names | pre_miRNA;  chromosome;  supercontig | - | | pre_miRNA;  chromosome;  supercontig;  biological_region;  SRP_RNA;  RNase_MRP_RNA | - | biological_region;  supercontig | | - | transposable_element;  transposable_element_gene;  chromosome | | - |
| Fasta file output | Yes | Yes | | Yes | Yes | Yes | | Yes | Yes | | Yes |
| Bed file output | Yes | No | | Yes | No | Yes | | No | Yes | | No |
| gff3/gtf file output | Yes | No | | Yes | No | Yes | | No | Yes | | No |
| Duplications | No | Yes | | No | Yes | No | | No | No | | No |
| Accuracy | High | Low | | High | Low | High | | Low | High | | Low |

Table 3: comparison between GAD and gff2sequence using GTF and FASTA files as inputs.

| Organism | *Drosophila melanogaster* | | *Oryza sativa* | | | | *Plasmodium malariae* | | | *Saccharomyces cerevisiae* | |
| --- | --- | --- | --- | --- | --- | --- | --- | --- | --- | --- | --- |
| Program/script | GAD | gff2sequence | | GAD | gff2sequence | GAD | | gff2sequence | GAD | | gff2sequence |
| Genes | 17753 | 1 | | 36844 | 31 | 6507 | | 33 | 7127 | | 40 |
| Gene SO names | gene | gene | | gene | gene | gene | | gene | gene | | gene |
| Transcripts | 34793 | 1 | | 43397 | 31 | 6507 | | 33 | 7127 | | 40 |
| Transcript SO names | transcript | transcript | | transcript | transcript | transcript | | transcript | transcript | | transcript |
| Up-stream | 17753 | 1 | | 36844 | - | 6507 | | - | 7127 | | - |
| Dwon-stream | 17753 | 1 | | 36844 | - | 6507 | | - | 7127 | | - |
| Intergenic regions | 18974 | - | | 37111 | 25 | 7926 | | 28 | 7298 | | - |
| 5`UTRs | 46320 | 1 | | 45538 | 29 | 337 | | 2 | 4 | | - |
| 3`UTRs | 33907 | 1 | | 51046 | 28 | - | | - | - | | - |
| Introns | 152701 | - | | 149091 | - | 7568 | | - | 380 | | - |
| Exons | 187494 | - | | 192488 | - | 14075 | | - | 7507 | | - |
| CDSs | 160948 | 1 | | 164924 | 22 | 13526 | | 21 | 6913 | | 1 |
| Other feature(s) SO names | start_codon;  Selenocysteine | - | | start_codon;  stop_codon | - | start_codon;  stop_codon | | - |  | |  |
| Fasta file output | Yes | Yes | | Yes | Yes | Yes | | Yes | Yes | | Yes |
| Bed file output | Yes | No | | Yes | No | Yes | | No | Yes | | No |
| gff3/gtf file output | Yes | No | | Yes | No | Yes | | No | Yes | | No |
| Duplications | No | No | | No | No | No | | No | No | | No |
| Accuracy | High | Low | | High | Low | High | | Low | High | | Low |
